# A Gene Therapy for Hereditary Nonpolyposis Colorectal Cancer using CRISPR-Cas9 Nickase

**DOI:** 10.1101/2023.06.20.545835

**Authors:** Shravan Kannan, Joshua J Man

## Abstract

Hereditary non-polyposis colorectal cancer (HNPCC) is an inherited disorder characterized by an increased risk of developing colorectal cancer before age 50. HNPCC is predominantly caused by genetic mutations in MLH1 and MSH2, which are involved in DNA mismatch repair. Current standard practice is to perform prophylactic colectomy, resulting in debilitating aftereffects for life. Though the genetic cause of HNPCC is well-known, there are currently no available treatments that target these mutations. Herein we describe a novel treatment protocol using a CRISPR-Cas9n-based genetic therapy to restore DNA mismatch repair. First, gRNA and template DNA targeting the most prevalent mutation clusters in MLH1 and MSH2 as well as CRISPR-Cas9n elements will be packaged into an integrase-deficient lentiviral vector. Then, the viral vector will be used to transduce human colonic tumor-derived organoids as well as administered systemically in mouse models of HNPCC. Mice will be monitored clinically and for signs of disease progression. At termination, colonic tissue will be harvested and analyzed for restoration of the wild-type MLH1 and MSH2 sequence and biochemical markers of HNPCC. This protocol offers an alternative strategy using CRISPR-Cas9n-based gene therapy to prevent tumor formation in patients, avoid morbid surgery, and significantly improve quality of life.

## INTRODUCTION

Lynch syndrome is a debilitating hereditary disorder impacting multiple organs, most commonly the colon, rectum, endometrium, prostate, and pancreas (**Idos and Valle, 2004**). Currently, there is no cure for Lynch syndrome, and the standard of care is to perform prophylactic colectomy in adolescence or chemotherapy and surgery in the event of tumor development. This syndrome is caused by mutations in genes involved in the DNA mismatch repair pathway, resulting in microsatellite instability and other tumorigenic mutations. Within Lynch syndrome, one of the most prevalent and debilitating diseases is hereditary non-polyposis colorectal cancer (HNPCC). While HNPCC is known to be caused by germline mutations, there are currently no genetic therapies available for HNPCC.

Due to the toxicities of traditional chemotherapy and the possibility of complications from surgical interventions, scientists are turning towards gene therapies to cure cancer. CRISPR (Clustered Regularly Interspaced Short Palindromic Repeats) are DNA sequences from foreign viral DNA that are integrated into the bacterial genome. When bacteria are re-infected, complementary RNA transcribed from CRISPR sequences can bind and induce Cas9-mediated cleavage, effectively providing anti-viral memory (**Barrangou, 2015**). *S. pyogenes* Cas9 (Cas9) has been re-engineered for gene editing directed by designer guide RNA (gRNA) containing a complementary sequence to the intended target. As an endonuclease, Cas9 cleaves double-stranded DNA three base pairs away from the protospacer adjacent motif (PAM) sequence (**Synthego, Full Stack Genome Engineering, 2022**). Double-stranded DNA breaks induce two major mechanisms for DNA repair – non-homologous end joining (NHEJ) and homology-directed repair (HDR) (**Tang et al**., **2019**). NHEJ merges the two blunt ends with DNA ligase. This is the most efficient DNA repair mechanism however at the expense of introducing indel mutations, which can lead to the deletion of important genetic material and frameshifts.

On the other hand, HDR can replace mutated DNA sequences at double-stranded DNA break sites with a wild-type sequence by introducing template DNA. This form of DNA repair is less efficient but more accurate. A substantial limitation of using the native Cas9 endonuclease is if double-stranded DNA breaks are executed at an unintended genomic locus and introduce off-target mutations. To mitigate this risk, researchers have modified the catalytic domain of Cas9 such that the resulting Cas9 “nickase” (Cas9n), induces only a single-stranded DNA break (**Trevino and Zhang, 2014**). Even if Cas9n creates a single-stranded DNA break in an unintended locus, it is unlikely that a second off-target single-stranded DNA break would occur in proximity to induce inappropriate DNA repair mechanisms. A sole single-stranded DNA break would be easily repaired by DNA ligase without introducing new DNA mutations.

This paper will describe a novel treatment method protocol for hereditary non-polyposis colorectal cancer (Lynch syndrome) that uses CRISPR-Cas9n to replace mutation clusters in MLH1 and MSH2 with their canonical sequences associated with normal health and intact DNA mismatch repair mechanisms. This gene therapy aids in preventing the development of colorectal cancer and eliminates the need for prophylactic colectomy in patients with Lynch syndrome. This theoretical therapy provides a framework for future genetic therapies responsible for Lynch syndrome that may fall outside of the area studied in the MLH1 and MSH2 genes.

## RESULTS

### Guide RNA Design and Validation

Mutations that result in loss of function or have a clinical association with HNPCC were mapped onto the protein structural domains of MLH1 and MSH2 (**Figure 1, adapted from Deihemi et al**., **2017**). The amino acid locations of the largest mutation clusters were in exon 2 of MLH1 and exon 12 of MSH2 which correspond to the ATPase binding domain of MLH1 and the MutL homolog interaction domain of MSH2, respectively. Potential 20-nucleotide gRNAs ending in the PAM sequence recognized by the SpCas9 protein were identified in the genomic sequences of MLH1 and MSH2 and assessed for predicted off-target binding. The final gRNAs I discovered when designing the treatment protocol were the following:

> **MLH1_Forward: 5’- CCCTCCATCGTAAATATAAA -3’**
>
> **MLH1_Reverse: 5’- GGATCAGGGTAAGTAAAACC -3’**
>
> **MSH2_Forward: 5’- CTTGATGAAAGGCCCAGTAT -3’**
>
> **MSH2_Reverse: 5’- CAGGTAAGTGCATCTCCTAG -3’**

**Figure 1:**
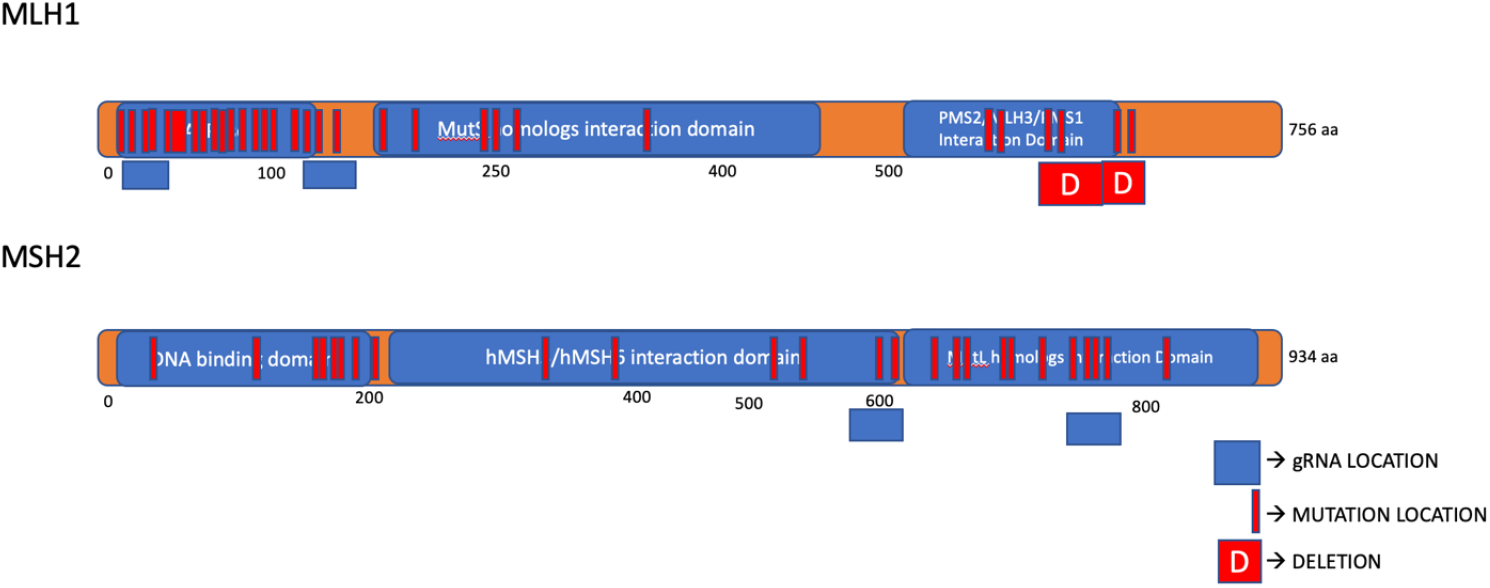
MLH1 and MSH2 Protein Structural Domains Annotated with Mutation Sites Reported in Humans. (adapted from Deihemi et al., 2017)

BLAST and CasOFFFinder were used to verify that these gRNAs specifically bind to the target sequence. BLAST predicted that these gRNAs do indeed bind to the intended genetic target sequence in humans. CasOFFFinder predicted possible off-target binding sites for gRNAs with up to 3 base pair mismatches as shown previously. (**Lin et al**., **2020**) gRNA sequences were accepted if they had at most 5 predicted off-target binding sites with 3 base pair mismatches and no off-target binding sites with 1 or 2 base pair mismatches compared to the target sequence. The results after verification of the gRNAs for off-target effects can be seen in Figures 4-7.

### TREATMENT PROTOCOL FOR HNPCC GENE THERAPY

#### Molecular Cloning of an Integrase-Deficient Lentiviral (IDLV) Packaging Plasmid Containing gRNA and SpCas9n

The IDLV packaging plasmid will be cloned as described previously (**Vijayraghavan and Kantor, 2017**). In brief, annealed oligos will be created using the 20-nucleotide gRNA sequences described above with the overhangs of the BbsI restriction enzyme sequence at the 5’ end according to the protocol by Addgene (*Addgene: Plasmid Modification by Annealed Oligo Cloning*). To combine Cas9 and gRNA elements into the same construct for lentiviral packaging, the pLentiCRISPR/v2 plasmid (Sanjana et al, 2014) will be digested with BbsI. Then, the annealed oligo pair with BbsI overhangs will be inserted and ligated. Next, the catalytic domain of Cas9 will be modified to make single-stranded DNA breaks as Cas9 nickase (Cas9n) by PCR-based site-directed mutagenesis as previously described (**Liu and Naismith, 2008**). Briefly, a pair of forward and reverse primers bearing a T->G point mutation will be used to introduce a D10A amino acid substitution, yielding Cas9n. BsrGI restriction enzymes will be used to digest the PCR product and create DNA overhangs to clone the Cas9n sequence into the pLentiCRISPR/v2 plasmid (**Figure 2A**).

**Figure 2:**
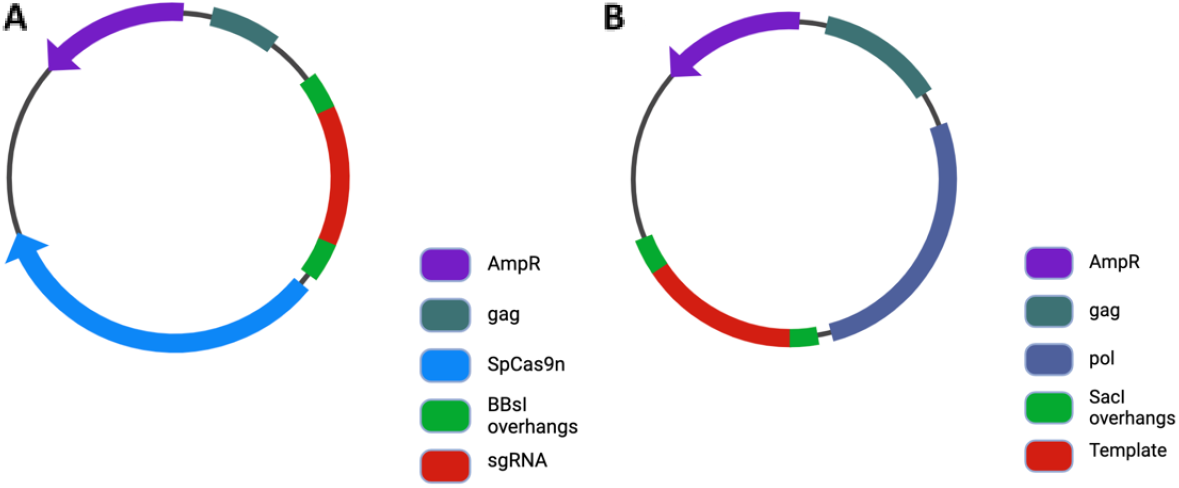
Map of the pLentiCRISPR/v2-gRNA (A) and psPAX2-HDRtemplate (B) plasmids. pLentiCRISPR/v2-gRNA and the psPAX2-HDRtemplate were cloned and digested with BbsI and the oligos were also digested with BbsI as well. These oligos were ligated as part of the plasmids.

#### Creation of a DNA Template for Homology-Directed Repair (HDR)

The nucleotide sequence flanked by the forward and reverse guide RNAs used to target exon 2 of MLH1 and exon 12 of MSH2 are 416 bp and 413 bp, respectively. Thus, a DNA template with homology arms of 200 bp before and after the genomic target sequence was designed to induce HDR, with a total length of 816 bp for MLH1 and 813 bp for MSH2. The wild-type genomic sequence was acquired from the UCSC Genome Browser and will be cloned using the forward and reverse primers below. In addition, Sacl restriction enzyme sites will be inserted at the 5’ and 3’ ends of the forward and reverse primers. The primers I discovered for this aspect of this treatment protocol are the following:

> **MLH1_HDR_Forward: 5’-AAAAAAGAGCTCCTGCCTGGCTAATTTTGTATT-3’**
>
> **MLH1_HDR_Reverse: 5’-AAAAAAGAGCTCAGAAGAGAATAGATTTTAATC-3’**
>
> **MSH2_HDR_Forward: 5’-AAAAAAGAACTCGAAAGATTTGACCATACTGA-3’**
>
> **MSH2_HDR_Reverse: 5’-AAAAAAGAGCTCTCCCTTGAAGATAGAAATGT-3’**

The HDR template sequence flanked by SacI restriction enzyme sites will be cloned from wild-type genomic DNA extracted from donor human peripheral blood mononuclear cells by the following PCR protocol. The annealing temperature is set within 5ºC from the melting temperature of both forward and reverse primers.

> Cycle 1:
>
> Denaturation: 96°C for 5 minutes
>
> Annealing: 67°C for 30 seconds
>
> Elongation: 72°C for 60 seconds
>
> Cycle 2-40:
>
> Denaturation: 96 °C for 30 seconds
>
> Annealing: 67 °C for 30 seconds
>
> Elongation: 72 °C for 60 seconds

Following SacI restriction enzyme digestion, the psPAX2 plasmid backbone (Addgene #12260) and PCR product insert will be ligated in a 1:10 molar ratio using DNA ligase. The resulting psPAX2-HDRtemplate (**Figure 2B**) plasmid will be transformed into competent E. coli. To confirm the transformation of bacteria, E. coli will be plated on agar mixed with ampicillin. DNA from bacteria will be purified by miniprep (Qiagen). To confirm accurate cloning, the empty psPAX2 plasmid backbone, and cloned psPAX2-HDRtemplate plasmid will be digested with Bsu36I (6301) and NheI (7714). The products will be run on a 1% agarose gel to assess restriction length fragments by gel electrophoresis. We expect that the cloned plasmid will yield a longer fragment (1.9 kb) compared to the empty plasmid backbone (1.4 kb) with the insertion of the HDR template. (**Figure 3**)

**Figure 3:**
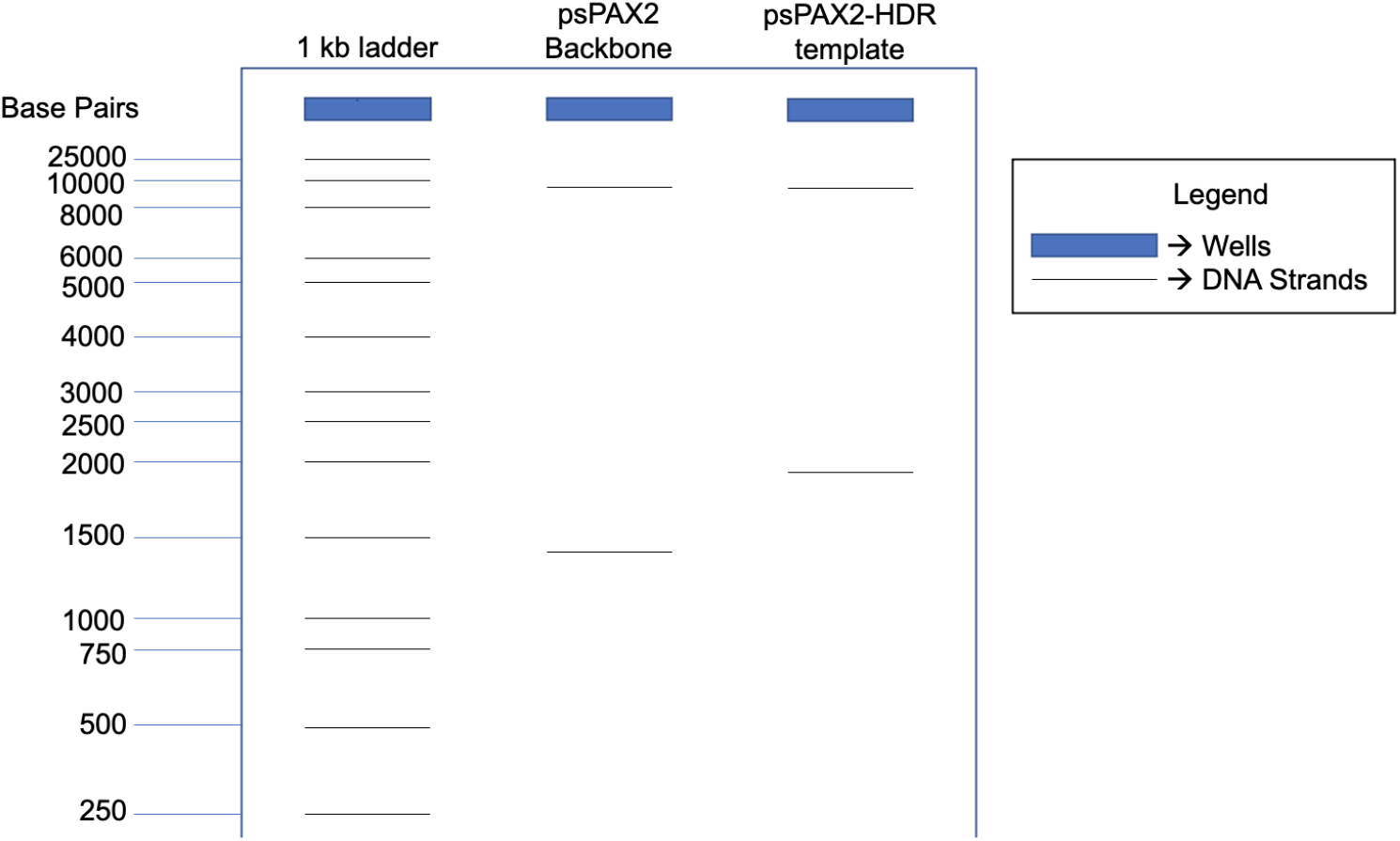
Expected gel results with psPAX2-backbone and psPAX2-HDRtemplate following digestion by Bsu36I (6301) and NheI (7714). psPAX2 plasmid backbone and psPAX2 HDR template will be run on 1% agarose gel for 2 hours. The blue rectangles represent wells and the lines represent the DNA strands.

**Figure 4:**
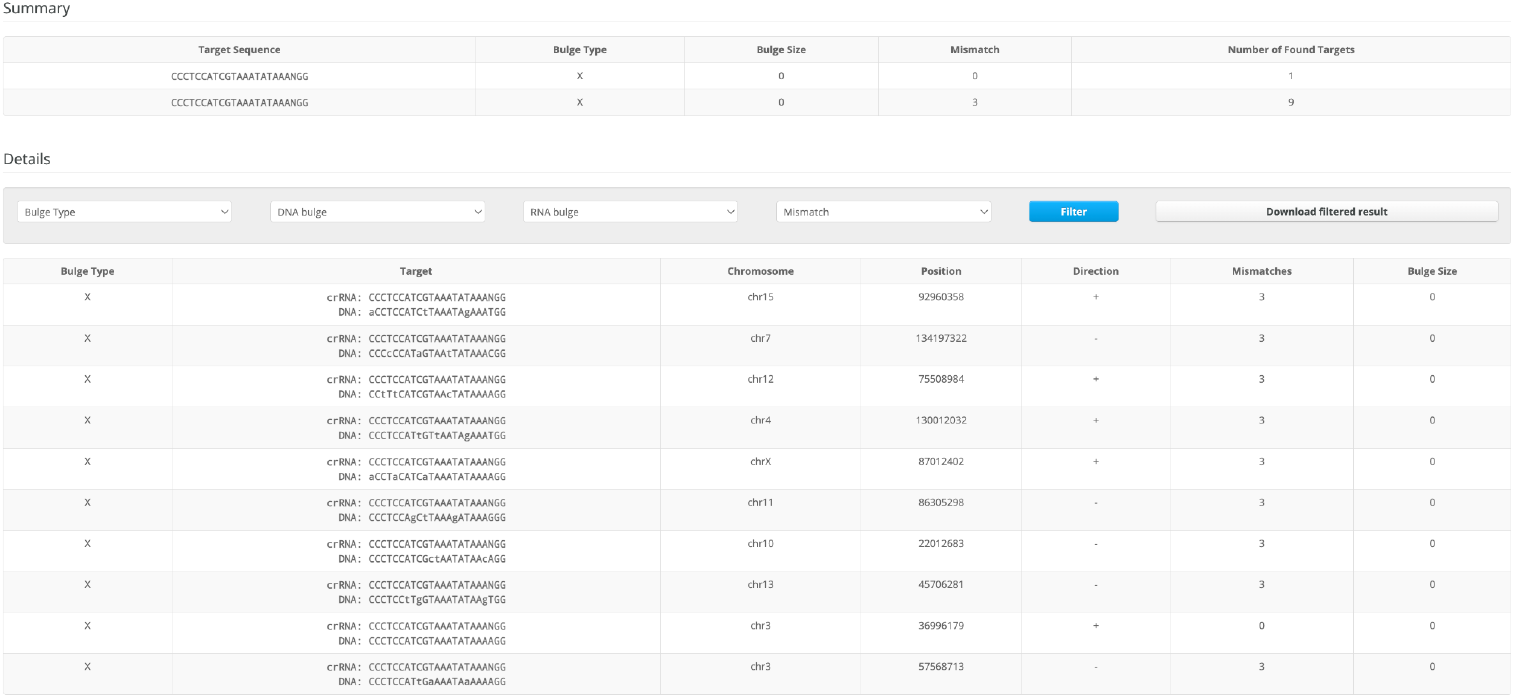
This is the result of gRNA verification done by CASOffFinder with 10 off-target effects for the forward gRNA of MLH1 in chromosomes 15, 7, 12, 4, 11, 10, 13, and 3.

**Figure 5:**
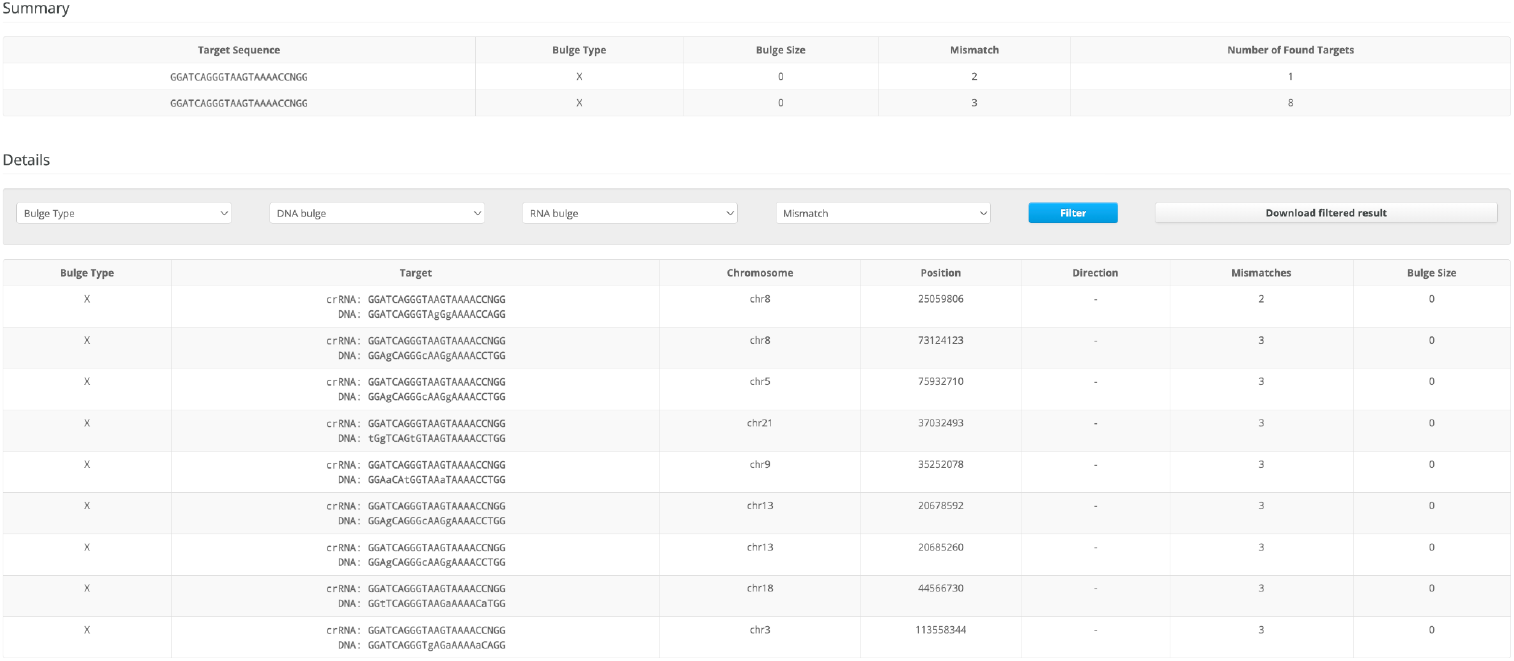
This is the result of gRNA verification done by CASOffFinder with 9 off-target effects for the reverse gRNA of MLH1 in chromosomes 8, 5, 21, 9, 13, 18, and 3.

**Figure 6:**
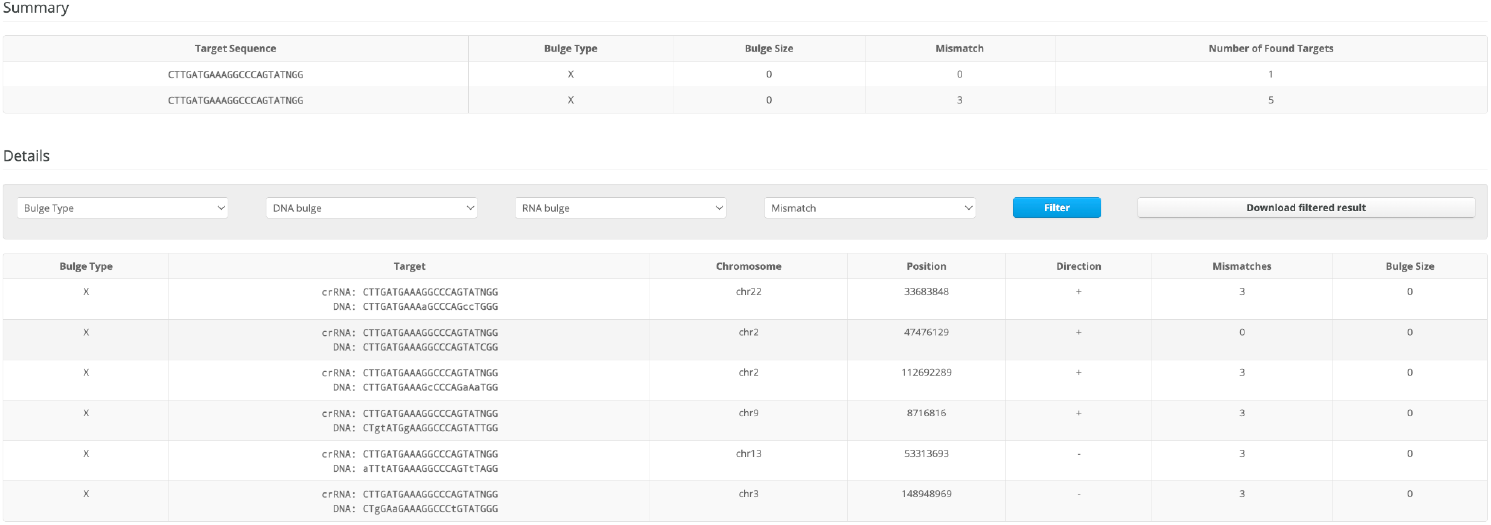
This is the result of gRNA verification done by CASOffFinder with 6 off-target effects for the forward gRNA of MSH2 in chromosomes 22, 2, 9, 13, and 3.

**Figure 7:**
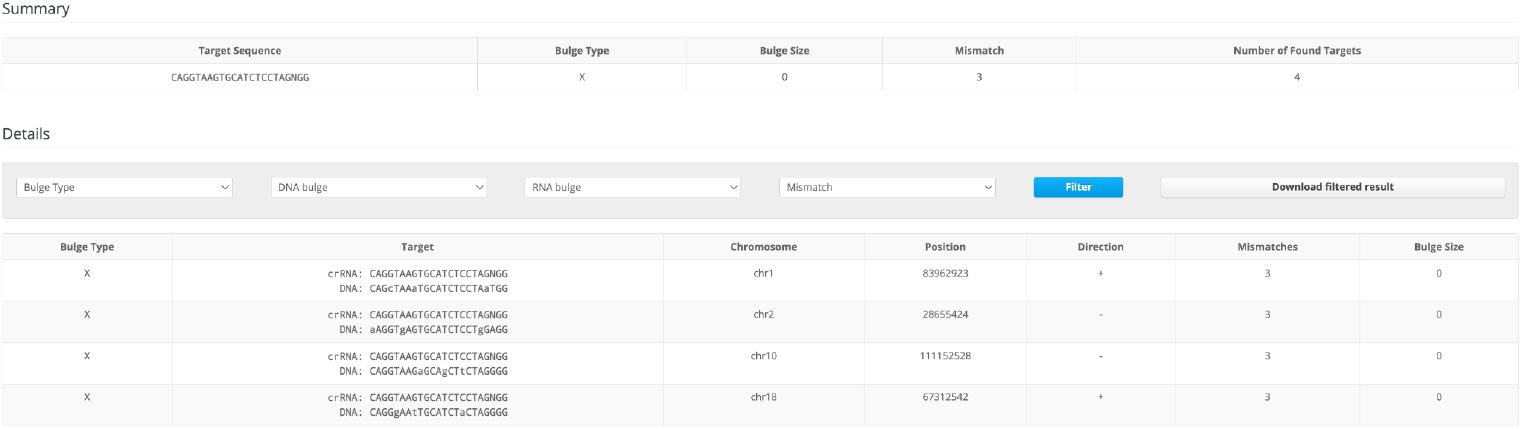
This is the result of gRNA verification done by CASOffFinder with 4 off-target effects for the reverse gRNA of MSH2 in chromosomes 1, 2, 10, and 18.

#### IDLV Packaging and Lentiviral Production

To package the gDNA, Cas9n, and HDR template elements into IDLVs, the pLentiCRISPR/v2-Cas9-gRNA and psPAX2-HDRtemplate plasmids will be transfected into HEK293 cells at 80% confluency with 1 M CaCl_2_ solution and a 2x BES-buffered solution as described previously (**Vijayraghavan and Kantor, 2017**). After 8 hours, HEK293 cells will be replenished with fresh media. Then after 24 hours, the supernatant will be transferred into separate tubes that will be ultracentrifuged and concentrated into pellets that will be resuspended in fresh media.

#### IDLV Transduction of Epithelial Cells from Human Colonic Explants

Viral concentration will be quantified by counting plaque-forming units (**Andersson and Lood. JoVE, 2019**) and adjusted to multiple particle concentrations that differ by a factor of 10. Patient-derived organoids (PDO) from HNPCC patients will be collected through colonoscopic polypectomies or prophylactic colectomies (**Okamoto et al., 2021**). To make conditioned media for PDOs, Noggin-expressing L1 cells will be grown in DMEM media until the cells reach 80% confluency. These cells will be incubated at 37ºC for 24 hours, and then the conditioned medium these cells are grown in will be collected. PDOs will be grown in a Noggin-conditioned medium for 3-6 days, changing the media every 3 days. The PDOs will be treated via injection of IDLVs containing pLentiCRISPR/v2-sgRNA and psPAX2-HDRtemplate versus the pLentiCRiPSR/v2 and psPAX2 backbones in the media. The viral transduction will be tested for multiple durations (3 days, 7 days, 14 days, and 21 days) with 1 injection per day to determine optimal transduction conditions.

#### Assessing Efficacy of Homology Directed Repair

To determine the efficacy of HDR-mediated restoration of the wild-type sequence, droplet-digital PCR (ddPCR) will be used to detect HDR or NHEJ events as described by the Gladstone Institute of Cardiovascular Disease (**Miyaoka et al, 2018**). ddPCR can divide up one PCR reaction into 20,000 water-in-oil droplets containing multiple copies of genome targets. These targets will be analyzed by HDR and NHEJ probes in each droplet. This method will be applied to genome DNA from colonic epithelia collected from the HNPCC mouse models below and HEK293 cells with knock-in exon 2 MLH1 mutations and exon 12 MLH2 mutations in vivo. The color coding provided by HDR, NHEJ, and reference probes in ddPCR will be used to determine the relative frequency of HDR versus NHEJ induced by IDLV treatment as described by Lambrescu et al. (2022).

#### Assessing Cellular and Morphologic Effects of IDLV Transduction

After viral transduction, the PDOs will be monitored for cellular and morphologic changes, including assays assessing cell survival, cell proliferation, and cell morphology. In cell survival assays, markers of apoptosis (e.g. Annexin V) and necrosis (e.g. miR-122, FK18, and HMCB1). will be quantified by Western blotting. In cell proliferation assays, cell count and phosphorylation of cyclin-dependent kinases will be assessed by microscopy and Western blotting. In cell morphology assays, the restoration of density-dependent inhibition will be assessed by microscopy before and after the injection of the treatment.

#### IDLV-Mediated Gene Therapy in a Murine Model of HNPCC

Mice-bearing mutations in exon 2 of hMLH1 (**Edelmann et al., 1996**) and exon 12 of hMSH2 (Jackson Labs #016231) will be obtained from Jackson Labs. 1000 nanograms (ng) of IDLV particles packaged with Cas9n, gRNAs, and the HDR template or IDLV particles packaged with just the pLentiCRISPR/v2 and psPAX2 backbones will be intrarectally administered into mice via lipophilic enemas (**Matsumoto et al., 2010**). The mice will then be monitored for changes in body weight, food consumption, stool composition, blood pressure, complete blood cell count, and electrolytes. The mice in each group will be treated for 21 days starting from 2 months old and sacrificed at 6 months old. The colonic epithelia will be harvested and assessed by H&E histology and immunohistochemical stains for carcinoembryonic antigens, carbohydrate antigens, tissue polypeptide specific antigens, and tumor-associated glycoprotein-72 (**Jelski and Mroczko, 2020**).

## DISCUSSION

In conclusion, we designed and validated gRNA that target MLH1 and MSH2, two genes involved in DNA mismatch repair that are commonly mutated in Lynch syndrome. We also present a treatment protocol to incorporate gRNA, Cas9 nickase, and a DNA template into IDLVs, transduce colonic epithelial crypt cells, and restore the canonical sequence of MLH1 and MSH2 by HDR.

We explained the use of CRISPR-Cas9n to introduce single-stranded DNA breaks in the loci of the highest density of MLH1 and MSH2 mutations reported in the literature that are associated with HNPCC or loss of mismatch repair function. As shown in **Figure 1**, these mutational clusters occur in exon 2 of MLH1 and exon 12 MSH2, which cause unique functional deficits. Mutations in exon 2 of MLH1 affect the ATPase binding domain and prevent micronuclei formation, thus detached chromosome fragments cannot be incorporated into nuclear DNA during cell division (**Jia and Chai, 2018**). Mutations in exon 12 of MSH2 affect the PMS2 interaction domain, which is required for the recruitment of MSH2 to sites of DNA mismatch to prevent microsatellite instability (**Li et al**., **2020**).

A major limitation that has precluded gene therapy from wider applications is the probability of off-target genetic editing that can lead to unintended novel mutations with potentially detrimental clinical side effects. When prior genomic editing approaches like zinc finger nucleases (ZFN) and transcription activator-like effector nucleases (TALEN) were used, it was difficult to predict off-target interactions between proteins and DNA. With RNA-DNA interactions, it is much easier to predict and minimize off-target binding; however, gRNA has been shown to bind to DNA with a tolerance of up to 3 base pair mismatches. Multiple methods were used in this theoretical experiment to reduce off-targeting. First, off-targeting sites were identified by using BLAST and CASOffFinder. Candidate gRNAs were ranked by prioritizing gRNA with the fewest single base pair interactions. Instead of using traditional Cas9, Cas9 nickase was used to introduce single-stranded breaks to minimize the probability of a double-stranded DNA break introducing unintended indel mutations by non-homologous end joining. Third, genomic DNA from colonic epithelial cells from mice or patient-derived organoids treated with IDLVs containing MLH1- and MSH2-targeting gRNA will be sequenced to assess for mutations at predicted off-target sites identified by CasOFFinder before clinical trials in patients. Thus, our protocol has highly specific gRNA sequences, a very low probability of inducing off-target mutations by NHEJ using Cas9n, and multiple means to assess safety in pre-clinical studies.

We also considered the possibility of off-targeting effects in other cell types besides colonic epithelial cells. The viral vector must not transduce non-proliferating cells like cardiac myocytes as this can cause serious toxicities for the patient, such as heart disease and cardiomyopathy. However, because germline mutations will affect the genome of every cell, non-specific transduction of other cell types, such as the pancreas and prostate, will also mitigate the risk of developing other cancers associated with Lynch syndrome.

To deliver the CRISPR-Cas9 elements efficiently and safely, the gRNAs need to be cloned into the psPAX2 packaging plasmid, which is separate from the integrase-deficient lentiviral (IDLV) packaging cassette and the VSV-G envelope plasmid. The genes encoding proteins necessary for viral replication (e.g. gag, pol, env, rev) are on separate plasmids to prevent unintended transduction of bacteria with new genes that could then be introduced into humans and result in potentially deleterious alterations. To prevent the genetic contents from integrating into the host genome, we specifically used a packaging plasmid with a modified int gene encoding a defunct integrase protein. IDLV was chosen because it cannot integrate its viral DNA elements into host cells, thus preventing unintended insertional mutations. Additionally, IDLVs do not induce lytic or lysogenic cycles. In addition, IDLVs have a reduced risk of mutations, compared to AAVs and lipid nanoparticles. AAVs have a lower immune response and a smaller packaging size of 4.7 kb. However, they have a lower carrying capacity and their immunotoxicity is dependent on tissues. On the other hand, lipid nanoparticles can be used to carry physical compounds, but are difficult to use. There is very little packaging size and nanoparticle-dependent immunotoxicity for lipid nanoparticles as well. The gene therapy also addresses several mutations clustered at foci spanning about 400 bp. Thus, an HDR template with flanking homology arms totaling over 800 bp was also required. Therefore, IDLVs were also favored for their larger packaging size. Additionally, transduction efficiency is expected to be higher for IDLVs compared with AAVs with a lower immunotoxicity compared to AAVs and lipid nanoparticles (**Biagioni et al**., **2018**).

To date, no gene therapies for HNPCC exist. A major challenge is the germline mutations driving the disease are present in all tissues, thus affecting DNA mismatch repair in all cells. However, the genetic treatment outlined in this paper addresses this issue. Although we propose to test our genetic therapy on the colonic epithelia, we envision that IDLVs can be administered systemically and thus target other tissues that are known to be at increased risk of carcinogenesis in Lynch syndrome. IDLVs also do not exhibit viral tropism, which is favored to non-specifically transduce virtually all cell types. However, this may be a disadvantage if high doses are needed to reach therapeutic levels in the colon. Future experiments will be performed to test whether HDR also restores the canonical sequence in tissues where cancers also arise (e.g. prostate, pancreas) and if concomitant intrarectal administration can achieve efficient gene editing of colonic crypt cells.

The proposed genetic therapy can potentially change the management of HNPCC dramatically. The current guideline for patients with HNPCC is to offer total colectomy at age 20 to prevent the development of colorectal cancer associated with high morbidity and mortality. The procedure is extremely invasive and can be associated with numerous post-operative morbidities (e.g. bleeding, infection, surgical adhesions, etc.). Most notably, this surgery forces patients to be dependent on parenteral nutrition at a young age for the rest of their life. The genetic therapy can avoid the need for colectomy by restoring the DNA mismatch repair pathway in colonic crypt cells, which is causally linked to colorectal oncogenesis in HNPCC. Longer-term studies of animal models of colorectal cancer in Lynch syndrome will help determine whether this gene therapy can prevent tumorigenesis.

The proposed genetic therapy has several limitations. This therapy would only be useful in patients with mutations in the specific foci in MLH1 and MSH2 targeted by our gRNA, thus this therapy is not indicated in patients with HNPCC in mutations in other domains of MLH1 and MSH2 or other MMR proteins. However, the general approach is universally applicable and can be specifically tailored to individual patients using the pipeline for newly designing appropriate gRNA described in this study. A second limitation is having multiple plasmids that need to be introduced to target cells via IDLVs, raising concerns about packaging efficiency. Lastly, high doses of IDLVs will likely be required for therapeutic efficacy. Whether intrarectal administration of IDLVs or another viral vector with tropism for colorectal crypt cells may allow for specific targeting at lower doses remains to be explored. Furthermore, IDLVs have never been assessed for genetic therapy in clinical trials. Thus, the safety profile of especially high doses of IDLVs needs to be clarified.

In conclusion, this protocol promotes the practice of personalized medicine. Although this treatment only targets two regions in two genes, it can target any region in those genes, as long as the gRNAs are designed with the rigorous checkpoints described in this study. This non-invasive approach addresses the underlying biological and molecular causes of Lynch syndrome compared to the current standard of care. Successful pre-clinical trials would raise the possibility of evaluating targeted genetic therapies in clinical trials in the future to assess their efficacy in improving patient outcomes and survival.

## METHODS

### Mapping Mutation Clusters in MLH1 and MSH2 in HNPCC

The human genomic sequences of MLH1 and MSH2 were acquired from the UCSC Genome Browser. Mutations in MLH1 and MSH2 that result in loss of function or have a clinical association with HNPCC were queried from the UniProtKB database (P40692) (P43246). These mutations were then mapped onto the structural domains of the MLH1 and MSH2 proteins to visually assess where mutational clusters occur and to inform guide RNA design.

### Guide RNA (gRNA) Design and Validation

Two guide RNAs (gRNA) flanking the ATPase binding domain of MLH1 and the MutL homolog interaction domain in MSH2 were designed to target the coding and complementary DNA strands. gRNAs were designed according to guidelines specific to the Streptococcus pyogenes Cas9 protein (SpCas9). (**Doench, 2020**) In brief, the genomic sequence was searched for available protospacer adjacent motif (PAM) sequences (e.g. 5’-NGG-3’ for the leading strand and a 5’-CCN-3’ for the lagging strand). The upstream 20 nucleotides were assessed for GC content. Off-target binding was assessed by Basic Local Alignment Search Tool (BLAST, NIH) and CasOFFFinder (Seoul National University).

## REFERENCES

Idos, Gregory, and Laura Valle. “Lynch Syndrome.” GeneReviews®, edited by Margaret P. Adam et al., University of Washington, Seattle, 1993, http://www.ncbi.nlm.nih.gov/books/NBK1211/.

Barrangou, Rodolphe. “The Roles of CRISPR–Cas Systems in Adaptive Immunity and Beyond.” Current Opinion in Immunology, vol. 32, Feb. 2015, pp. 36–41, doi:10.1016/j.coi.2014.12.008.

Synthego | Full Stack Genome Engineering. https://www.synthego.com/guide/how-to-use-crispr/pam-sequence. Accessed 28 July 2022.

Trevino, Alexandro E., and Feng Zhang. “Genome Editing Using Cas9 Nickases.” Methods in Enzymology, vol. 546, 2014, pp. 161–74, doi:10.1016/B978-0-12-801185-0.00008-8.

Deihimi, Safoora, et al. “BRCA2, EGFR, and NTRK Mutations in Mismatch Repair-Deficient Colorectal Cancers with MSH2 or MLH1 Mutations.” Oncotarget, vol. 8, no. 25, May 2017, pp. 39945–62, doi:10.18632/oncotarget.18098.

Addgene: Plasmid Modification by Annealed Oligo Cloning By. https://www.addgene.org/protocols/annealed-oligo-cloning/?gclid=Cj0KCQjwhY-aBhCUARIsALNIC05wzNFo3W5N1H9m3VR-8SElLJLijOxnHM6TXjAiyDIbYRp0fb-HG5MaAulaEALw_wcB. Accessed 11 Oct. 2022.

Sanjana, Neville E., et al. “Improved Vectors and Genome-Wide Libraries for CRISPR Screening.” Nature Methods, vol. 11, no. 8, 2014, pp. 783–84, doi:10.1038/nmeth.3047.

Liu, Huanting, and James H. Naismith. “An Efficient One-Step Site-Directed Deletion, Insertion, Single and Multiple-Site Plasmid Mutagenesis Protocol.” BMC Biotechnology, vol. 8, no. 1, Dec. 2008, p. 91, doi:10.1186/1472-6750-8-91.

Addgene: PsPAX2. https://www.addgene.org/12260/. Accessed 10 May 2023.

Vijayraghavan, Sriram, and Boris Kantor. “A Protocol for the Production of Integrase-Deficient Lentiviral Vectors for CRISPR/Cas9-Mediated Gene Knockout in Dividing Cells.” Journal of Visualized Experiments : JoVE, no. 130, Dec. 2017, p. 56915, doi:10.3791/56915.

Andersson, Tilde, and Rolf Lood. Determining Viral Titer as Plaque Forming Units (PFU) | Microbiology | JoVE. https://www.jove.com/v/10514/plaque-assay-method-to-determine-viral-titer-as-plaque-forming-u nits. Accessed 9 May 2023.

Okamoto, Takuya, et al. “A Protocol for Efficient CRISPR-Cas9-Mediated Knock-in in Colorectal Cancer Patient-Derived Organoids.” STAR Protocols, vol. 2, no. 4, Sept. 2021, p. 100780, doi:10.1016/j.xpro.2021.100780.

Miyaoka, Yuichiro, et al. “Detection and Quantification of HDR and NHEJ Induced by Genome Editing at Endogenous Gene Loci Using Droplet Digital PCR.” Methods in Molecular Biology (Clifton, N.J.), vol. 1768, 2018, pp. 349–62, doi:10.1007/978-1-4939-7778-9_20.

Lambrescu, Ioana, et al. “Application of Droplet Digital PCR Technology in Muscular Dystrophies Research.” International Journal of Molecular Sciences, vol. 23, no. 9, Jan. 2022, p. 4802, doi:10.3390/ijms23094802.

Edelmann, Winfried, et al. “Meiotic Pachytene Arrest in MLH1-Deficient Mice.” Cell, vol. 85, no. 7, June 1996, pp. 1125–34, doi:10.1016/S0092-8674(00)81312-4.

016231 - Strain Details. https://www.jax.org/strain/016231. Accessed 11 Oct. 2022.

Jelski, Wojciech, and Barbara Mroczko. “Biochemical Markers of Colorectal Cancer – Present and Future.” Cancer Management and Research, vol. 12, June 2020, pp. 4789–97, doi:10.2147/CMAR.S253369.

Jia, Pingping, and Weihang Chai. “The MLH1 ATPase Domain Is Needed for Suppressing Aberrant Formation of Interstitial Telomeric Sequences.” DNA Repair, vol. 65, May 2018, pp. 20–25, doi:10.1016/j.dnarep.2018.03.002.

Li, Juyi, et al. “A Novel Splice-Site Mutation in MSH2 Is Associated With the Development of Lynch Syndrome.” Frontiers in Oncology, vol. 10, 2020, https://www.frontiersin.org/articles/10.3389/fonc.2020.00983.

Biagioni, Alessio, et al. “Delivery Systems of CRISPR/Cas9-Based Cancer Gene Therapy.” Journal of Biological Engineering, vol. 12, Dec. 2018, p. 33, doi:10.1186/s13036-018-0127-2.

UniProt. https://www.uniprot.org/uniprotkb/P40692/entry. Accessed 10 May 2023.

UniProt. https://www.uniprot.org/uniprotkb/P43246/entry. Accessed 10 May 2023.

BLAST: Basic Local Alignment Search Tool. https://blast.ncbi.nlm.nih.gov/Blast.cgi. Accessed 10 May 2023.

CRISPR RGEN Tools. http://www.rgenome.net/cas-offinder/. Accessed 10 May 2023.

